# Distinct Gene Regulatory Dynamics Drive Skeletogenic Cell Fate Convergence During Vertebrate Embryogenesis

**DOI:** 10.1101/2024.03.26.586769

**Authors:** Menghan Wang, Ana Di Pietro-Torres, Christian Feregrino, Maëva Luxey, Chloé Moreau, Sabrina Fischer, Antoine Fages, Patrick Tschopp

**Affiliations:** DUW Zoology, University of Basel, 4051 Basel, Switzerland

**Keywords:** cell type evolution, cell fate convergence, vertebrate skeletogenesis, gene regulatory evolution, single-cell functional genomics

## Abstract

Cell type repertoires have expanded extensively in metazoan animals, with some clade-specific cells being paramount to their evolutionary success. A prime example are the skeletogenic cells of vertebrates that form the basis of their developing endoskeletons. Depending on anatomical location, these cells originate from three different embryonic precursor lineages – the neural crest, the somites, and the lateral plate mesoderm – yet they converge developmentally towards similar cellular phenotypes. Furthermore, these lineages have gained ‘skeletogenic competency’ at distinct timepoints during vertebrate evolution, thus questioning to what extent different parts of the vertebrate skeleton rely on truly homologous cell types.

Here, we investigate how lineage-specific molecular properties of the three precursor pools are integrated at the gene regulatory level, to allow for phenotypic convergence towards a skeletogenic cell fate. Using single-cell transcriptomics and chromatin accessibility profiling along the precursor-to-skeletogenic cell continuum, we examine the gene regulatory dynamics associated with this cell fate convergence. We find that distinct transcription factor profiles are inherited from the three precursor states, and that lineage-specific enhancer elements integrate these different inputs at the *cis*-regulatory level, to execute a core skeletogenic program.

We propose a lineage-specific gene regulatory logic for skeletogenic convergence from three embryonic precursor pools. Early skeletal cells in different body parts thus share only a partial ‘deep homology’. This regulatory uncoupling may render them amenable to individualized selection, to help to define distinct morphologies and biomaterial properties in the different parts of the vertebrate skeleton.

## Introduction

Metazoan bodies are defined by obligate multicellularity and the presence of functionally, morphologically, and molecularly distinct cell types^1-3^ On a developmental timescale, this requires the implementation of distinct gene regulatory programs, to generate cellular diversity from a single fertilized cell, the zygote, that inherits a common genomic blueprint to its cellular descendants^4-6^. Such regulatory programs can also inform us about evolutionary relationships amongst different cell types, both within and across species boundaries. Namely, on an evolutionary timescale, novel cell types are often thought to arise through the genomic re-use of pre-existing regulatory modules, *via* re-shuffling, neo-functionalization or co-option. Accordingly, similarities or dissimilarities in the gene regularity dynamics underlying their developmental specification can help to resolve potential cell type homologies^1,7^.

In this context, vertebrate skeletogenesis offers a relevant opportunity to study the gene regulatory dynamics of cell fate specification across both developmental and evolutionary timescales. A cartilaginous and often ossified endoskeleton is a hallmark of the vertebrate clade, with many accessory connective tissue types required to build a functional skeleton^8^. Developmentally, the specification of the cells required to build these tissues initiates with mesenchymal precursors condensing, at the onset of vertebrate skeletogenesis, and then progresses through distinct, cell type-specific differentiation processes^9-12^. A peculiarity in this process is that – unlike as for many other cell lineages (Fig.1A, left) – these multipotent skeletal progenitors can arise convergently, from distinct embryonic sources (Fig.1A, right). Namely, depending on anatomical location, three distinct embryonic lineages – the cranial neural crest, the somitic sclerotome, and the somatopleure of the lateral plate mesoderm – are the developmental origin of the cranial, axial and appendicular skeleton, respectively^8^ (Fig. 1B). The early specification of these cells thus needs to integrate discrete molecular states, inherited from their respective embryonic precursor sources, to facilitate a skeletogenic cell fate convergence.

**Figure 1.**
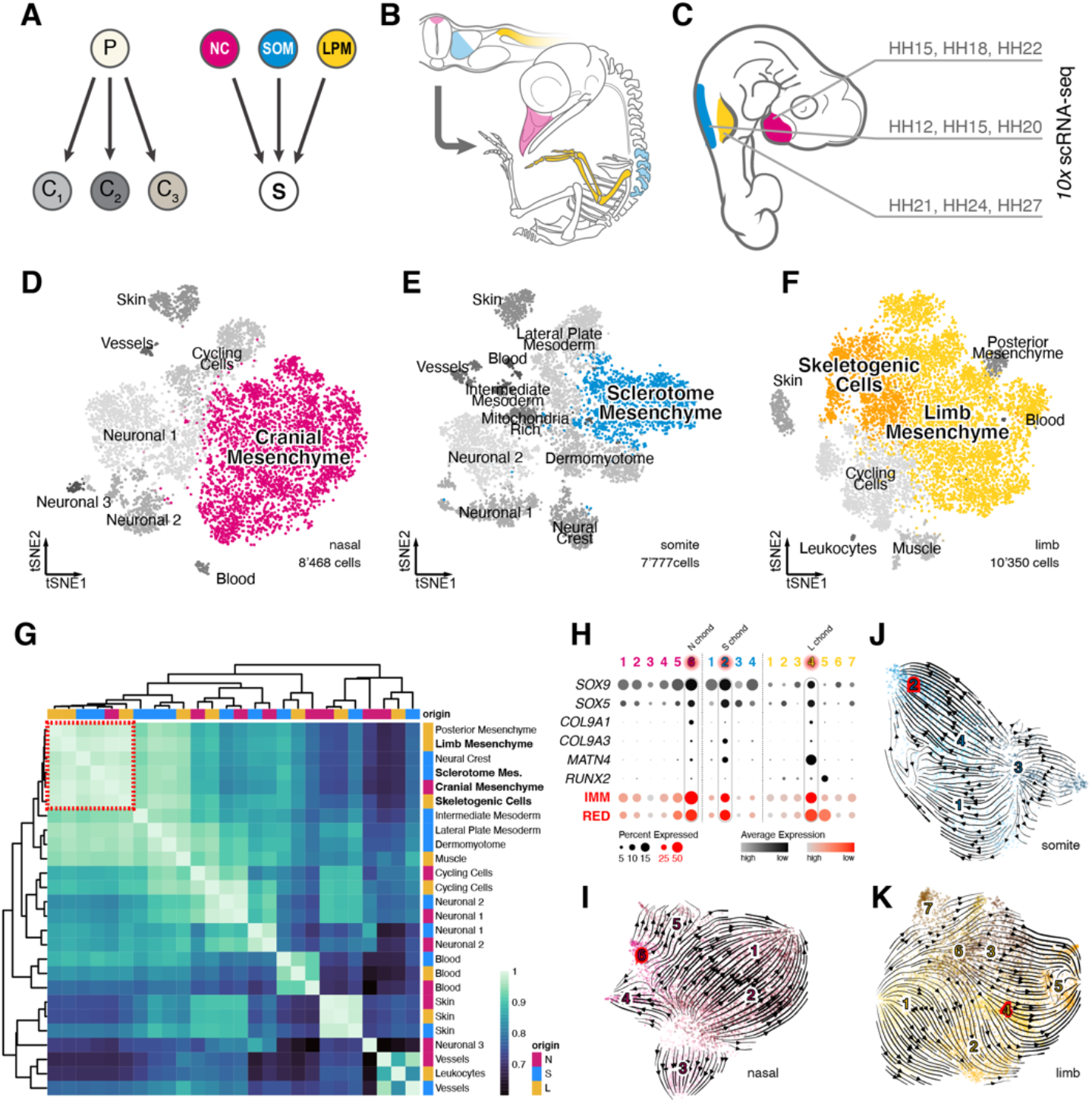
A convergent transcriptomic signature in skeletogenic cells of different embryonic origins. (**A**) Divergent diversifying *versus* convergent cell fate decisions. Precursor cells (P) usually differentiate into functionally distinct cell types (C1-3). During vertebrate skeletogenesis, however, three distinct embryonic lineages – the neural crest (NC, magenta), the somitic mesoderm (SOM, blue) and the lateral plate mesoderm (LPM, yellow) – give rise to functionally similar skeletogenic cells (S). (**B**) Early embryonic origins and eventual anatomical locations of skeletogenic cells sampled in this study. (**C**) Sampling scheme to assess the transcriptional dynamics of skeletogenic convergence across anatomical locations (color coded) and embryonic stages (Hamburger-Hamilton, HH). (**D**-**F**) tSNE representations of the three scRNA-seq datasets with ‘broad’ cell type annotations, with the mesenchymal populations used for ‘fine’ re-clustering highlighted in color. (**G**) Unsupervised hierarchical clustering and heatmap representation of pairwise Spearman’s rank correlation coefficients of highly variably genes in pseudobulk transcriptomes of ‘broad’ cell type clusters. Anatomical origins of pseudobulks are indicated by color code. Mesenchymal cell populations across embryonic origins cluster together (red dotted square). (**H**-**K**) ‘Fine’ re-clustering of mesenchymal populations. (**H**) Dot plot of chondrogenic marker genes (black) and chondrogenic modules (red) expression in ‘fine’ clusters identified across the three embryonic origins. (**I**-**K**) tSNE representations of re-clustered mesenchymal cells of nasal (**I**), somite (**J**) and limb (**K**) origins, with ‘fine’ cluster annotations and superimposed streamline plots of *scVelo* vector fields. Early chondrogenic clusters are highlighted in red.

The distinct embryonic trajectories of skeletogenic cells are also reflecting the evolutionary histories of the different parts of the vertebrate skeleton. The first vertebrate skeletal elements are thought to have originated in the head region, mirroring the prevertebrate presence of cartilage-like structures supporting a feeding apparatus^13^. Subsequently, the ability to form a progenitor cell-based endoskeleton expanded along the primary and secondary body axes, giving rise to structurally supportive yet flexible elements in the axial and appendicular skeletons, respectively^14^. However, these distinct evolutionary and developmental histories may also challenge our understanding to what extent the convergently specified cells of the vertebrate skeleton can be considered truly homologous^15-17^.

Here, using single-cell functional genomics along the three distinct mesenchymal precursor-to-skeletogenic cell trajectories, we investigate the genome-wide regulatory dynamics in a vertebrate embryo at cellular resolution. We provide evidence that lineage-specific transcription factor profiles are inherited from the respective embryonic origins and that these are integrated at distinct *cis*-regulatory elements, to canalize developmental cell fate trajectories towards an early skeletogenic convergence point. These distinct dynamics imply a regulatory uncoupling between skeletogenic cells at different anatomical locations. We discuss the resulting implications for cell type homology assessment in the vertebrate skeleton, and the potential of distinct evolutionary trajectories in skeletal cell and tissue properties, upon co-option-dependent convergence of gene regulatory programs across embryonic lineages.

## Results

### A convergent transcriptomic signature in skeletogenic cells of different embryonic origins

To establish the temporal progression of skeletogenic initiation and maturation across the three anatomical locations and embryonic lineages, we first performed chromogenic *in situ* hybridizations (ISH) on a developmental time series of chicken embryos. We used cranial sagittal cryosections covering the frontonasal prominence, and brachial trunk transversal sections including the emerging forelimb buds. We investigated the expression of *SOX9*, an early marker of skeletogenic induction^9^, and *Aggrecan* (*ACAN*), an extracellular matrix protein of more mature skeletal cells^18^ (Fig.S1A,B). Based on the observed expression dynamics, we devised a sampling strategy following the cell- and tissue-specific transcriptional changes of the three embryonic lineages, from mesenchymal precursors towards the onset and maturation of skeletogenic tissues, using *10x Chromium* single-cell RNA-sequencing (scRNA-seq) profiling (Fig.1C). In total, we obtained over 23’000 high quality single-cell transcriptomes. For each of the three anatomical locations, we integrated the three sampled embryonic stages and performed tSNE non-linear dimensionality reduction and graph-based clustering using *Seurat*^19^ (Fig.1D-F). These ‘broad’ clusters were annotated with the help of expression profiles of known marker genes (Fig.S1C,E). All samples showed comparable transcriptome complexities, and contributed similarly to the different clusters (Fig.S1F,G).

To assess transcriptomic similarities amongst these clusters, we next generated cluster-based pseudobulks and calculated Spearman’s rank correlation coefficients on differentially expressed genes across all cell types and anatomical locations. Unsupervised hierarchical clustering revealed that cell types originating from the same embryonic lineage, but sampled at different anatomical locations – like, e.g., skin or blood cells – showed highly similar transcriptional profiles, in agreement with their shared developmental history (Fig.1G). Intriguingly, however, our analysis revealed that also mesenchymal cells stemming from discrete embryonic lineages – that is, from the neural crest, the somites, or the lateral plate mesoderm – clustered together, indicating a transcriptional convergence amongst them (Fig.1G, red dotted square). Across our three anatomical sampling sites, we focused on ‘broad’ clusters that likely contained cells transitioning from a mesenchymal precursor state towards a skeletogenic fate (Fig.1D-F, color coded), and re-clustered them at finer resolutions. Within these ‘fine’ clusters, we again used expression profiles of known marker genes, as well as two previously identified early chondrogenic gene co-expression modules, ‘IMM’^20^ and ‘RED’^21^, to assess their respective skeletogenic differentiation. In each embryonic lineage, we identified a ‘fine’ cluster enriched for an early skeletogenic, i.e. chondrogenic, signature (Fig.1H, highlighted in red). For the limb sample, we additionally isolated a cluster showing signs of more mature skeletal cells (Fig.1H, limb ‘fine’ cluster 5). Using *scVelo*^22^, we approximated cell fate transition trajectories *in silico* and projected the predicted vector fields using streamline plots on tSNE representations of our mesenchymal samples. For all three anatomical locations, *scVelo* predicted trajectories connecting the mesenchymal precursors – as identified by known marker genes – to early chondrogenic populations (Fig.1I-K).

Using ISH and single-cell transcriptomics, our analyses revealed distinct temporal dynamics, but converging molecular signatures amongst mesenchymal cells with skeletogenic potential across the three embryonic lineages. Furthermore, our *scVelo* analysis suggested that our single-cell transcriptomics data captured the entire specification spectrum, from uncommitted mesenchymal precursor cell to early chondrocyte.

### Distinct *trans*- and *cis*-regulatory modalities underlie the convergent specification of skeletogenic cells

To detail the transcriptional signatures underlying the switch from uncommitted mesenchymal precursor cell to early chondrocyte, we focused our analyses on the ‘fine’ mesenchymal clusters with skeletogenic potential. We re-assessed transcriptional similarities of these sub-populations using differentially expressed genes and Spearman’s rank correlation coefficients of pseudobulk transcriptomes. Again, we found high transcriptional similarities amongst early skeletogenic cells at the three anatomical locations, as indicated by them clustering together (Fig.2A, red dotted square).

**Figure 2.**
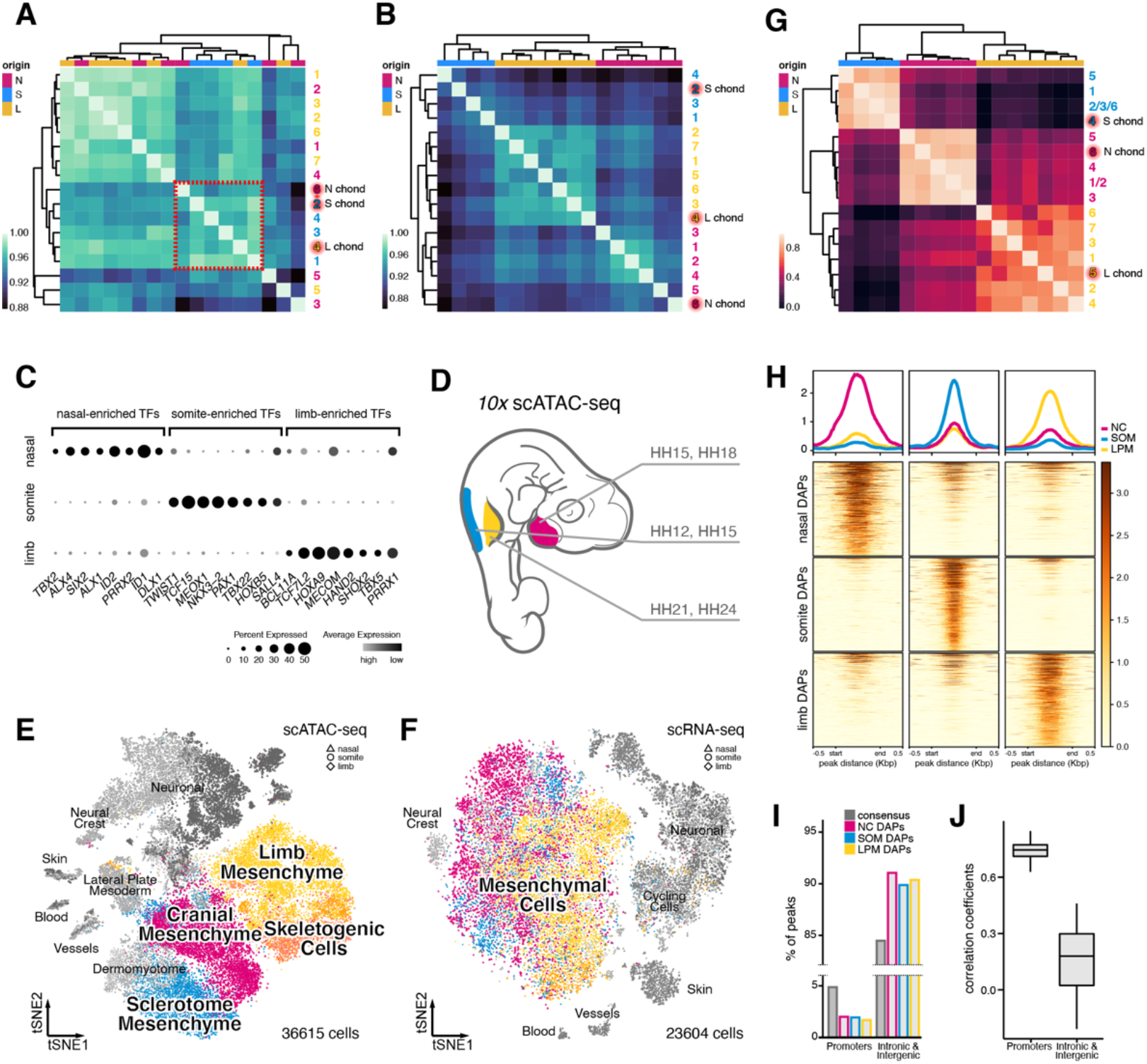
Distinct *trans*- and *cis*-regulatory modalities underlie the convergent specification of skeletogenic cells. (**A**) Unsupervised hierarchical clustering and heatmap representation of pairwise Spearman’s rank correlation coefficients of highly variably genes in pseudobulk transcriptomes of ‘fine’ mesenchymal clusters. Anatomical origins of pseudobulks are indicated by color code. Early chondrogenic cells (highlighted in red) across embryonic origins cluster together (red dotted square). (**B**) Unsupervised hierarchical clustering and heatmap representation of pairwise Spearman’s rank correlation coefficients of expressed transcription factors in pseudobulk transcriptomes of ‘fine’ mesenchymal clusters. Anatomical origins of pseudobulks are indicated by color code. All pseudobulks, including early chondrogenic cells (highlighted in red), cluster by embryonic origins. (**C**) Dot plot of embryonic origin-enriched transcription factors expression in early chondrogenic cells. (**D**) Sampling scheme to assess chromatin accessibility signatures of skeletogenic cells across anatomical locations (color coded) and embryonic stages (Hamburger-Hamilton, HH). (**E**-**F**) tSNE representations of integrated single-cell chromatin accessibility (**E**) and single-cell transcriptome (**F**) data across the three anatomical locations. Anatomical origins of cells are indicated by symbols, with the mesenchymal populations used for ‘fine’ reclustering highlighted in color. (**G**) Unsupervised hierarchical clustering and heatmap representation of pairwise Spearman’s rank correlation coefficients of differentially accessible peaks in pseudobulk chromatin accessibility data of ‘fine’ mesenchymal clusters. Anatomical origins of pseudobulks are indicated by color code. All pseudobulks, including early chondrogenic cells (highlighted in red), cluster by embryonic origins. (**H**) Coverage plots and heatmap representations of the top500 differentially accessible peaks (DAPs) in early chondrogenic cells of neural crest, somitic and lateral plate mesoderm origin. (**I**) Promoter depletion and intronic/intergenic elements enrichment of DAPs in early chondrogenic cells, relative to the consensus peak set. (**J**) Boxplots of pairwise Spearman’s rank correlation coefficients of DAPs in early chondrogenic cells across embryonic origins, calculated using promotor-proximal elements or an equal number of randomly sampled distal intronic/intergenic elements.

To investigate potential upstream regulatory inputs controlling these similar transcriptomes, we next focused our attention on transcription factors. Spearman’s rank correlation coefficients on transcription factor expression profiles, however, re-clustered the mesenchymal sub-populations strictly by embryonic origins, irrespective of their skeletogenic differentiation state (Fig.2B). This suggested that the mesenchymal precursor populations carried over a lineage-specific repertoire of expressed transcription factors, while undergoing skeletogenic induction. Indeed, when looking at transcriptional regulators enriched in chondrogenic cells of the three anatomical locations, we find clear evidence of a lineage-specific heritage of expressed transcription factors, many of which have known developmental functions in their respective anatomical locations (Fig.2C). Thus, counterintuitively, this indicated that – across anatomical locations and embryonic lineages – overall similar transcriptional signatures were generated with distinct upstream *trans*-regulatory inputs, i.e. lineage-specific transcription factor expression profiles.

We reasoned that these distinct *trans*-regulatory inputs could potentially be integrated at the *cis*-regulatory level of key skeletogenic genes, to facilitate transcriptomic convergence. To investigate this possibility, we performed single-cell chromatin accessibility assays (single-cell assay for transposase-accessible chromatin with sequencing, or scATAC-seq^23^) to identify genomic elements with potential regulatory activity. We followed a similar sampling scheme as for our scRNA-seq approach, although – reasoning that such *cis*-regulatory recoding would occur during early stages of skeletogenic induction – we excluded the late time points (Fig.2D). In total, we obtained over 36’000 cells with high quality chromatin accessibility profiles and, using *MACS2*^24^, we identified a consensus set of 678,707 peaks across the three anatomical locations. We integrated the two embryonic stages per sampling site, performed non-linear dimensionality reduction and identified potential cell type-specific clusters (Fig.S2A,C). We annotated these clusters with the help of our scRNA-seq data, using label transferring and non-negative least squares (NNLS) regression, as well as visual inspection of scATAC-seq ‘marker peaks’ (see Material and Methods, and Fig.S2D). At all three anatomical locations, overall similar cell type repertoires were recovered as in our scRNA-seq sampling. We then combined all samples across the three anatomical locations, for both scRNA-seq and scATAC-seq data sets, and performed anchor-based integration. Mesenchymal cells in our scATAC-seq data visually appeared to intermingle less than in our scRNA-seq data (Fig.2E,F). This may indicate the presence of embryonic origin-specific states in chromatin accessibility, as opposed to the convergent signatures at the transcriptomic level (Fig.1G). To investigate potential lineage-specific chromatin accessibilities, we re-clustered the mesenchymal scATAC-seq cell populations with skeletogenic potential and annotated the resulting ‘fine’ clusters using our scRNA-seq data (Fig.S2E,G). We then performed unsupervised hierarchical clustering on Spearman’s rank correlation coefficients of differentially accessible peaks (DAPs) in pseudobulks of these ‘fine’ mesenchymal clusters. Akin to our scRNA-seq analyses of transcription factor expression profiles, this resulted in a strict embryonic origin-dependent clustering of mesenchymal populations, including early chondrogenic cells (Fig.2B,G). Major cell type classifications, however, still clustered according to their ‘broad’ annotations, with similar chromatin accessibility signatures echoing their shared embryonic origins (Fig.S2H). This further implied that mesenchymal and skeletogenic cells across the three anatomical locations carry distinct chromatin accessibility profiles, reflecting their discrete embryonic origins and lineage histories.

Indeed, coverage plots across the three anatomical locations revealed that a substantial fraction of DAPs within the respective chondrogenic populations are distinct and hence embryonic origin-specific (Fig.2H). We classified these DAPs according to their genomic location, with respect to the transcription start sites of neighboring genes. Interestingly, we found that promoter-proximal peaks were depleted in our DAP set, relative to the consensus gene set, while more distal elements – intronic and intergenic – appeared enriched (Fig.2I). This implied that the majority of differences in chromatin accessibilities, between chondrogenic cells from different embryonic lineages, originated at distally located peaks. Indeed, DAPs at promoter-proximal peaks showed on average higher Spearman’s rank correlation coefficients across embryonic origins than distal intronic/intergenic ones (Fig.2J). This suggested that the similar transcriptional profiles observed at the RNA level originate from similar promoter repertoires. Distal peaks, however, where one would anticipate putative long-range enhancer elements to be located, showed much lower similarity in accessibilities amongst the different embryonic origins (Fig.2J).

Collectively, our combined scRNA-seq and scATAC-seq approach revealed the presence of distinct *trans*- and *cis*-regulatory signatures in skeletogenic cells at the three anatomical locations, as evidenced by embryonic origin-specific transcription factor expression profiles and the presence of discrete chromatin accessibility signatures at distal locations.

### *Trans*- and *cis*-regulatory dynamics of skeletogenic convergence across the three embryonic lineages

To follow the *trans*- and *cis*-regulatory changes underlying these convergent cell fate transitions, we next investigated scRNA-seq and scATAC-seq dynamics along chondrogenic pseudotime trajectories in the three anatomical locations. Using tSNE embeddings of our scRNA-seq data, we constructed minimum spanning trees using *slingshot*^25^, with preset start points corresponding to the clusters with naïve mesenchymal expression signatures (Fig.3A-C). We observed overall similar trajectory predictions as for our *scVelo* analysis (Fig.1H-J). For each anatomical location, we were able to retrieve a single trajectory either ending in an early chondrogenic cluster (Fig.3A,B), or traversing it towards more mature skeletal cells (Fig.3C). We transferred the pseudotime values of our chondrogenic scRNA-seq trajectories to our scATAC-seq data, to follow the accompanying chromatin accessibility dynamics (Fig.3A-C, insets).

**Figure 3.**
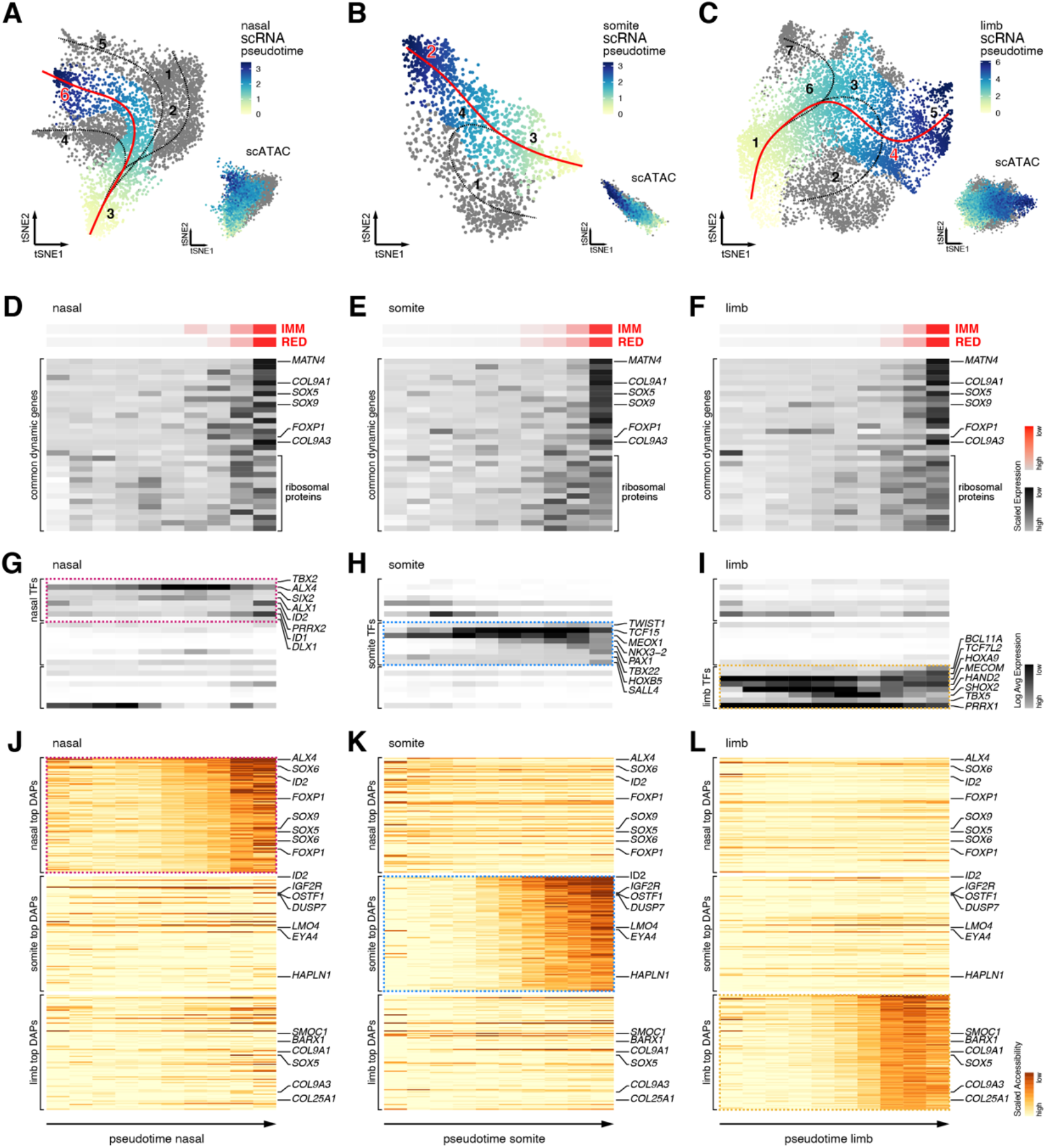
*Trans*- and *cis*-regulatory dynamics of skeletogenic convergence across three embryonic lineages. (**A**-**C**) tSNE representations of single-cell transcriptomes and co-embedded single-cell chromatin accessibilities (insets) for nasal (**A**), somite (**B**), and limb (**C**) origins. Superimposed on the single-cell transcriptomes are the pseudotime trajectories identified by *slingshot*, with the chondrogenic trajectories used for further analyses highlighted in red. Pseudotime progression visualized by heatmaps on scRNA and scATAC data. ‘Fine’ cluster annotations as in Figure 1, with early chondrogenic clusters highlighted in red. (**D**-**L**) Binned pseudobulk dynamics along chondrogenic pseudotime trajectories. (**D**-**F**) Z-score scaled expression dynamics along the chondrogenic pseudotime trajectories for chondrogenic modules (red) and common differentially expressed genes identified in all three embryonic origins (black). (**G**-**I**) Log-transformed expression dynamics of embryonic origin-specific transcription factors identified in nasal (magenta dotted line), somite (blue dotted line), and limb (yellow dotted line) samples. (**J**-**L**) Average normalized peak accessibility dynamics of embryonic origin-specific differentially accessible peaks (DAPs) identified in nasal (magenta dotted line), somite (blue dotted line), and limb (yellow dotted line) samples. Select DAP-adjacent genes are indicated on the right.

We then binned the respective scRNA-seq pseudotimes into equidistant pseudobulks and used *TrAGEDy*^26^ to align the chondrogenic gene expression dynamics along the respective trajectories of the three embryonic origins. We found overall higher similarities towards the ends of these pair-wise comparisons, indicating chondrogenic convergence at the transcriptional level (Fig.S3A,C). Both nasal and somite trajectories failed to align to the last section of our limb chondrogenic pseudotime. This corroborated the notion that we had recovered more mature skeletal cells in our limb samples, compared to nasal and somite (Fig.S3A,F, Fig.1H). Accordingly, we excluded the corresponding limb pseudotime bins containing these cells from further analyses.

We first checked the expression dynamics of the two chondrogenic gene co-expression modules ‘IMM’ and ‘RED’. Both modules increased in their expression along the three pseudotime trajectories, in nasal, somite and limb samples (Fig.3D-F, red). Next, we looked at the expression dynamics of a core set of common chondrogenic genes, found to be enriched in chondrocytes across the three embryonic lineages (Fig.S4A). All genes showed an increase in expression along the respective trajectories (Fig.3D-F, black). Contained within this shared set of genes were known chondrogenic regulators and extracellular matrix proteins, as well as a selection of ribosomal proteins (Fig.3D-F, Fig.S4A). Additionally, using *tradeSeq*^27^, we identified lineage-specific expression dynamics of known and novel regulators activated in chondrogenesis across the three anatomical locations (Fig.S4B,D). In all three lineages, the chondrogenic wave appeared to be preceded by an increase in expression of origin-specific transcription factors (Fig.3G-I, Fig.2C). Finally, we found evidence for distinct chromatin accessibility dynamics. Chondrocyte-specific DAPs showed increased accessibilities along the respective chondrogenic scATAC pseudotime trajectories, but in an embryonic origin-specific manner (Fig.3J-L).

Using integrative scRNA-seq and scATAC-seq pseudotime analyses, we detailed the emergence of common transcriptional signatures and distinct *trans*- and *cis*-regulatory profiles during skeletogenic convergence. Namely, while increased expression of chondrogenic modules and a core set of differentially expressed genes was shared across the three embryonic origins, these were accompanied by lineage-specific transcription factor expression and chromatin accessibility dynamics.

### Embryonic origin-specific activities and specificities of transcription factor binding motifs

To investigate the interplay of distinct transcription factor profiles and origin-specific chromatin accessibilities, we next evaluated cell type-specific activities of transcription factor binding motifs. Given the scarcity of publicly available and experimentally validated binding motifs for chicken transcription factors, we decided to define our own set of DNA position weight matrices. Briefly, we used *Homer*^28^ to identify enriched *de novo* motifs in our scATAC-seq data in a cluster-by-cluster manner, across the three anatomically distinct samples. We annotated these *de novo* motifs with candidate transcription factors using public repositories, and selected the best matches based on motif similarity and the correlation of motif activities with scRNA-seq transcription factors expression profiles (see Material and Methods). In total, we identified and annotated 1373 *de novo* motifs across the three anatomical locations (Fig.S5A), with 540 non-redundant ones used for the further analyses (Fig.4A). Motifs for members of the homeobox, C2H2 zinc fingers and basic helix-loop-helix (bHLH) protein transcription factor families were amongst the most frequently identified ones (Fig.4A). Furthermore, our limb mesenchyme *de novo* motif for SOX9 matched a chick limb ChIP-seq^29^ validated motif more closely than publicly available position weight matrices (Fig.S5B). Encouraged by this, we continued all subsequent analyses with our *de novo* motifs only.

**Figure 4.**
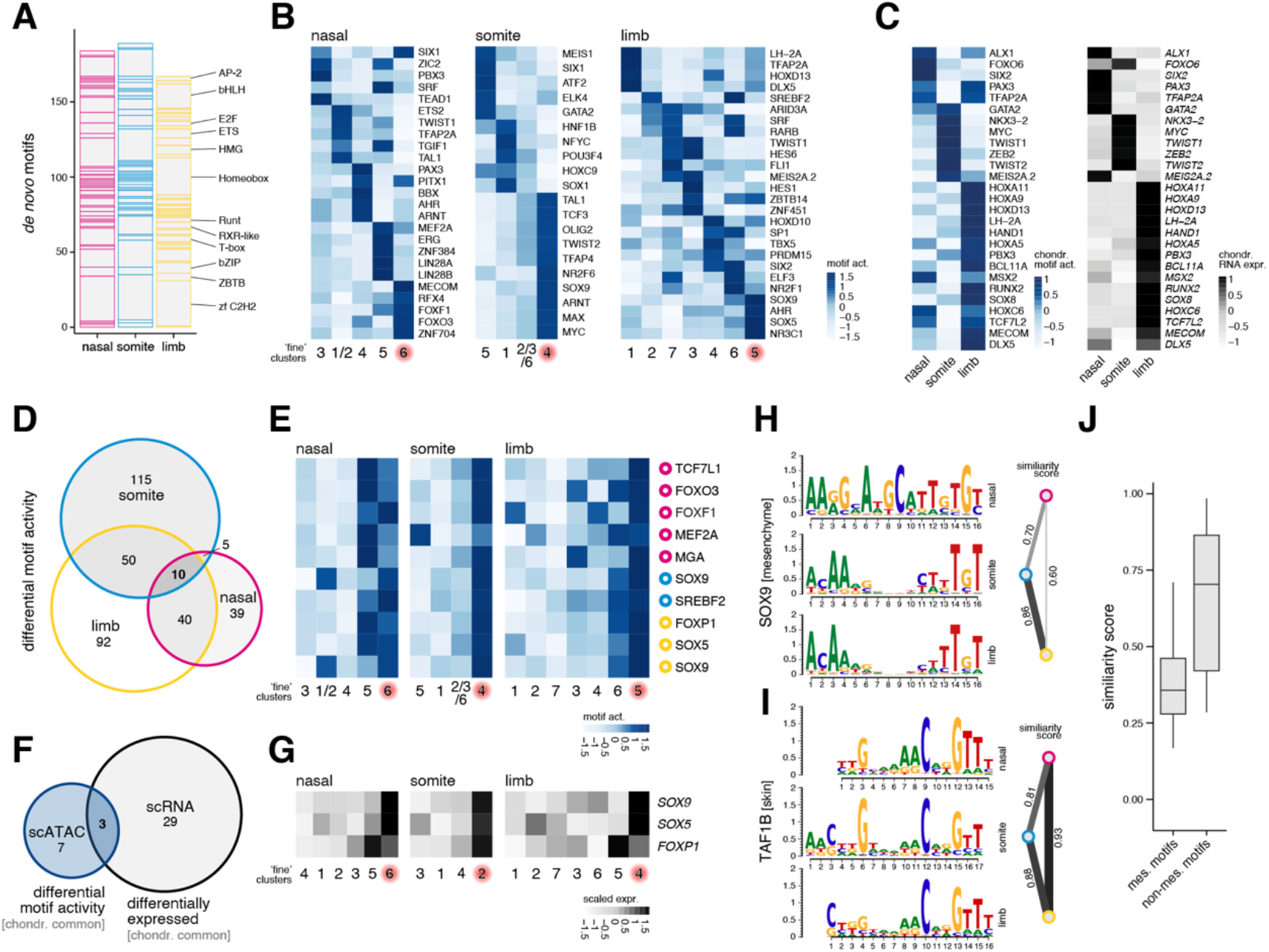
Embryonic origin-specific *de novo* prediction and activity profiles of transcription factor binding motifs. (**A**) Final numbers and transcription factor (TF) family distribution of non-redundant *de novo* binding motifs across the three embryonic origins. (**B**) Differential cluster-specific motif activities in mesenchymal populations across embryonic origins. ‘Early chondrogenic’ clusters are highlighted in red. (**C**) Embryonic origin-specific TF motif activities (left) and mRNA expression profiles (right), enriched in early chondrogenic cells. (**D**) Venn diagram displaying the overlap of chondrocyte-enriched motif activities across embryonic origins. (**E**) Motif activity heatmaps of the ten commonly chondrocyte-enriched TF motifs across embryonic origins. The embryonic origin in which the respective motif was identified is indicated by color-coded circles on the right. ‘Early chondrogenic’ clusters are highlighted in red. (**F**) Venn diagram displaying the overlap of unique chondrocyte-enriched motif activities and commonly enriched chondrogenic genes. (**G**) Enriched expression of *SOX9, SOX5, FOXP1* in early chondrogenic cells of the three embryonic origins. (**H, I**) Position weight matrices of a TF motif identified in mesenchymal cells of all embryonic origins (*SOX9*, **H**) and a TF motif identified in non-mesenchymal cells (skin) of all embryonic origins (*TAF1B*, **I**). Pairwise motif similarity scores are displayed on the right, with wider/darker lines indicating higher similarity. (**J**) Boxplots of pairwise motif similarity scores for TFs identified in multiple embryonic origins. Motifs were binned according to which general cell population they were identified in, i.e. mesenchymal *versus* non-mesenchymal.

Using this custom set of position weight matrices, we next conducted differential motif activity analyses. At both ‘broad’ and ‘fine’ cluster resolutions, we identified motif activity signatures that were predictive for specific cell types, including early chondrocytes and mesenchymal precursor cells (Fig.4B and Fig.S5C,D). Moreover, we identified motif activities in chondrogenic cells with high embryonic origin-specificity. These were mirrored by expression signatures of the corresponding transcription factors. This emphasized the presence of distinct *trans*-regulatory inputs in chondrogenic cells of the three embryonic lineages, at both RNA and motif activity levels (Fig.4C). Of all motifs whose activities were enriched in chondrogenic cells, only ten were shared across the three anatomical locations (Fig.4D,E, Fig.S5E). Comparing them to the core chondrogenic genes identified in our differential expression analyses revealed only three genes showing consistent chondrocyte enrichment at both RNA expression and motif activity levels: *SOX9, SOX5* and *FOXP1* (Fig.4F,G).

With the length range of our *Homer de novo* motif search (8-22 bp) we were able to investigate the occurrence of potential co-binding patterns of multiple transcription factors. Sequences bound by multimeric protein complexes are increasingly recognized as an integral aspect of DNA’s regulatory grammar, to provide robustness or diversify activity patterns in a combinatorial manner^30-32^. Indeed, many of our motifs showed a bimodal distribution of nucleotide enrichment in their position weight matrices, indicative of dimers binding to them. For example, the SOX9 motifs identified in somite and limb mesenchymal cells indicated a homodimer-like binding, in agreement with previous findings^29^ (Fig.4H and Fig.S5B). Additionally, in nasal samples, our analyses predicted a motif showing a SOX9-like monomer signature at its 3’ end, but a potential heterodimeric binding partner at its 5’ end (Fig.4H). To investigate this phenomenon more systematically – that is, the same transcription factor having dissimilar binding motifs predicted in the three different embryonic lineages – we trimmed the extremities of our position weight matrices based on minimal nucleotide enrichment scores and calculated motifs similarities across anatomical locations. In our analysis, we split the motifs based on whether their position weight matrices were identified in mesenchymal cells of different embryonic origins, or in non-mesenchymal populations of the same embryonic lineage (e.g. skin, Fig.4I). On average, position weight matrices for transcription factors identified in mesenchymal cells showed lower similarity scores than motifs identified in non-mesenchymal cell types (Fig.4J). This indicated that binding motifs for the same transcription factors were less conserved in mesenchymal cells of different embryonic origins, potentially due to lineage-specific differences in their cells’ chromatin environment, or distinct co-factors binding the motifs in a heterodimeric manner (Fig.4H-J).

Collectively, we identified *de novo* transcription factor binding motifs, some of which showed common cell type-specific activities while others were embryonic lineage-restricted. Furthermore, we find partially diverging sequence logos for the same transcription factors in mesenchymal cells at different anatomical locations. This substantiates our previous findings of lineage-specific *trans*-regulatory inputs during the skeletogenic convergence of different embryonic precursor lineages at the level of motif architectures and activities.

### Lineage-specific regulatory architectures and enhancer activities in skeletogenic cells

We used *ArchR*^33^ on our mesenchymal scATAC-seq data for peak-to-gene links analyses, to connect putative distal enhancer elements to target genes and test for lineage-specific activities. Based on a minimal correlation coefficient and adjusted p-value cutoffs, we identified over 28’000 presumptive peak-to-gene links (Fig.5A-C and Fig.S6A,C). Within these peak-to-gene links, target genes were generally predicted to be contacted by less than three putative *cis*-regulatory elements (CRE), and the majority of CREs was interacting with only one target gene (Fig.S6D,F). Using hierarchical k-means (hkmeans) clustering, we sorted our peak-to-gene links according to their aggregate activity profiles and plotted z-score normalized chromatin accessibilities and imputed target gene expression levels (Fig.5A-C and Fig.S6G,I). For both somite and limb samples, and to some extent for nasal cells, we identified clusters enriched for early chondrogenic cells (Fig.5A-C), with corresponding enrichments for skeletogenesis-related terms (Fig.S6G,I).

**Figure 5.**
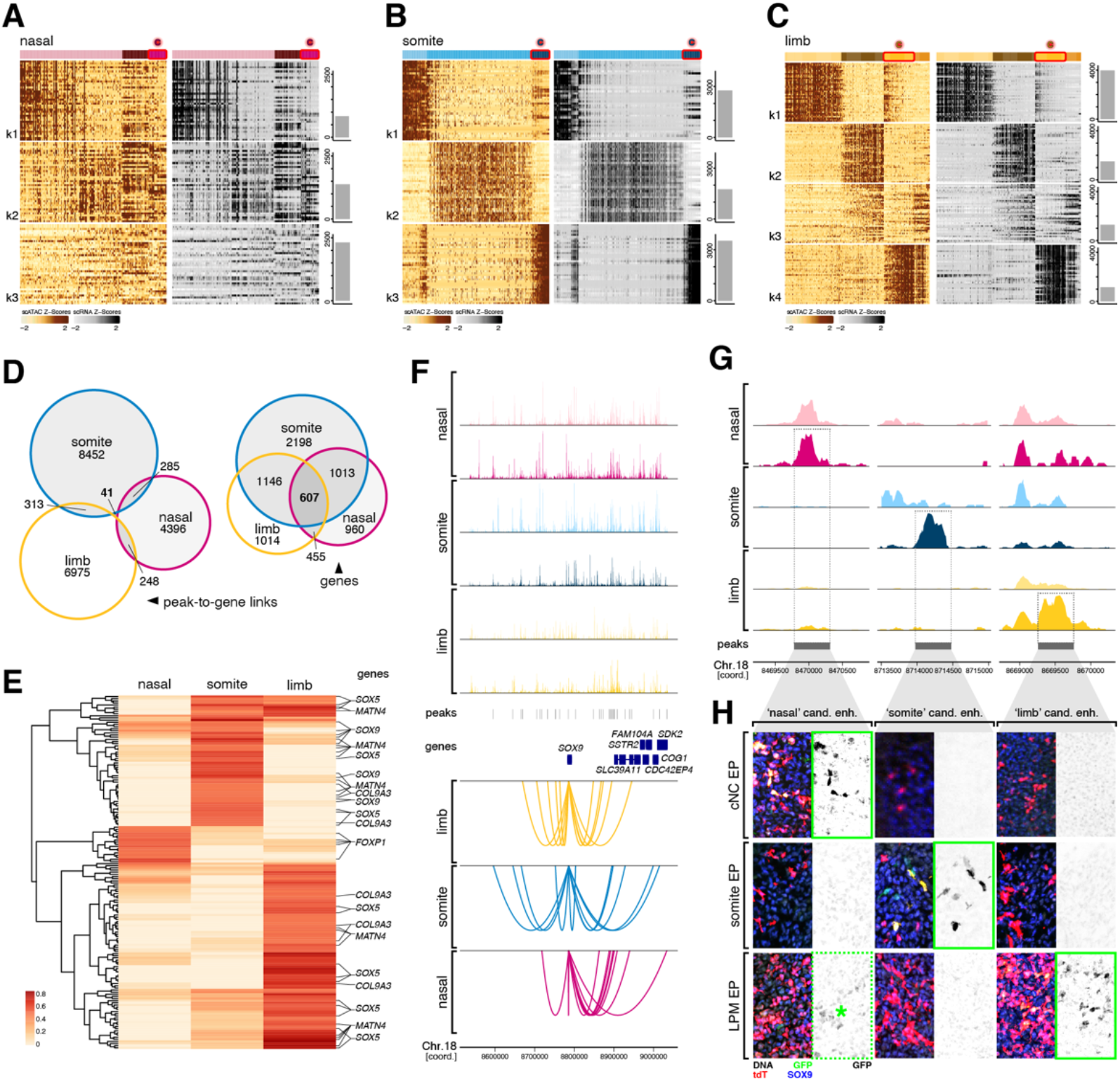
Lineage-specific regulatory architectures and enhancer activities in skeletogenic cells. (**A**-**C**) hkmeans-clustered peak-to-gene link heatmaps displaying Z-score normalized peak accessibilities (left, ochre) and imputed target gene expression levels (right, grey) in single cells (columns). Cell clusters dominated by early chondrogenic cells are highlighted in red (top). The top40 peak-to-gene links are shown. Total numbers of identified peak-to-gene links per cluster are indicated by barplots on the right, cluster numbers (kX) on the left. (**D**) Venn diagram displaying the overlap of mesenchymal peak-to-gene link CREs across embryonic origins (left) and overlap of target genes contained within these peak-to-gene links (right). (**E**) Hierarchically clustered peak-to-gene link correlation heatmap of links identified at core chondrogenic genes, across embryonic origins. Select target genes are indicated on the right. (**F**) Pseudobulk aggregate genome accessibility tracks at the *SOX9* locus. For each embryonic origin, tracks for precursor populations (light colors) and early chondrogenic cells (dark colors) are displayed. Identified peaks and neighboring genes are indicated. Below, embryonic origin-specific peak-to-gene link plots. (**G**) Pseudobulk aggregate genome accessibility tracks of peaks tested in enhancer reporter assays in **H**. Coordinates of candidate enhancers are highlighted by grey dotted boxes. Precursor population tracks in light colors, early chondrogenic tracks in dark colors. (**H**) *In vivo* enhancer reporter assays. Chondrocyte candidate enhancer elements with predicted embryonic origin-specificity were cloned into reporter constructs driving GFP expression and electroporated (EP) together with a tdTomato control plasmid into cells of the cranical neural crest (cNC), the somites, or the forelimb lateral plate mesoderm (LPM). Embryonic origin-specificity of GFP expression in SOX9-positive chondrogenic condensations is indicated by green squares. The candidate ‘nasal’ enhancer showed weak GFP signal in the limb mesenchyme as well (green dotted square, asterisk).

Of all peak-to-gene links, only 41 CRE coordinates were detected in all three anatomical locations using our stringent cutoffs. However, that number increased to 607, if we were only focusing on the overlap of predicted target genes therein (Fig.5D). This suggested that many commonly expressed genes were contacted by lineage-specific enhancer elements. Indeed, looking at the correlation scores of our core chondrogenic gene set (Fig.S4A, excluding ribosomal proteins) revealed largely lineage-specific peak-to-gene link activities (Fig.5E).

We further explored this in more detail at the *SOX9* locus, a well-known regulator of skeletogenic induction^9^. While a high density of consensus peaks was present in the genomic landscape flanking that gene, the overall pseudobulk chromatin accessibility profiles looked distinct from one anatomical location to another, both in mesenchymal precursors (bright colors) as well as at the early chondrogenic stage (dark colors) (Fig.5F, top). Importantly, link plots at the locus revealed that many of the peak-to-gene links were in fact predicted to be embryonic origin-specific (Fig.5F, bottom). To evaluate putative enhancer functions of these peaks and test for their lineage-specificity, we first used differential chromatin accessibility analysis to define chondrocyte-enriched DAPs at the *SOX9* locus. For all three anatomical locations, we identified peaks with high chromatin accessibilities in early chondrogenic cells, relative to mesenchymal precursors and chondrocytes of other embryonic origins (Fig.5G). We isolated the corresponding sequences from genomic DNA and cloned them into reporter plasmids, upstream of a minimal promoter driving green fluorescent protein (GFP) expression. We electroporated the resulting constructs into the precursor populations of the respective skeletogenic lineages *in ovo*. As electroporation control, we included a plasmid containing a strong constitutively active promoter driving tdTomato expression. Post-electroporation, we let the embryos develop further for two days, harvested the targeted tissues and processed them for histology. We performed immunohistochemistry against endogenous SOX9 protein, to determine the location of chondrogenic condensations, and against tdTomato and GFP, to evaluate electroporation efficiency and enhancer activity, respectively. Within electroporated condensations, we scored the presence or absence of GFP signal, to indicate chondrogenic enhancer activity. The predicted somite and limb enhancers showed specific GFP reporter activity, restricted to the respective embryonic lineages they were identified in (Fig.5H, green boxes). The nasal candidate enhancer reporter drove strong GFP signal in cranial neural crest-derived chondrogenic tissue (Fig.5H, green box). It additionally did so at reduced levels in limb condensations as well (Fig.5H, green dotted box and asterisk), suggesting only partial lineage-specificity of this element.

Overall, our peak-to-gene link analyses and enhancer reporter assays uncovered the presence of lineage-specific CREs contacting an overlapping set of target genes across the three anatomical locations. This suggested a partial lineage dependency in *cis*-regulatory interactions of target genes in early chondrogenic cells of different embryonic origins, mirroring their previously established specificity at the *trans*-regulatory level.

## Discussion

Cell type specification in animals relies on the execution of distinct gene regulatory programs during embryonic and post-embryonic development. Consequently, cell type evolution depends on the origination of new regulatory modalities, to drive innovations in cellular form and function. Here we have documented the gene regulatory dynamics underlying the embryonic specification of skeletogenic cells in vertebrates, an iconic cell type central to their evolutionary success. Following their first evolutionary appearance at the base of vertebrates, additional embryonic lineages have subsequently acquired the ability to form skeletal cell types. This makes the underlying gene regulatory programs interesting from both evolutionary and developmental perspectives. Namely, how the three embryonic lineages have acquired skeletogenic competency at a gene regulatory level during vertebrate evolution, and, developmentally, how these lineages are transcriptionally recoded during the earliest steps of skeletogenesis, to converge from distinct molecular profiles of their origin populations towards a similar skeletogenic phenotype (Fig.1A).

### Transcriptional recoding and lineage-memory in mesenchymal cells

Using scRNA-seq profiling, we have followed the early transcriptional dynamics of vertebrate skeletogenesis in three distinct embryonic lineages, at three different anatomical locations: cranial neural crest (frontonasal prominence), somitic sclerotome (axial skeleton) and lateral plate mesoderm (appendicular skeleton) (Fig.1B). We find shared transcriptional signatures between both mesenchymal and skeletal cells of the three embryonic origins (Fig.1G). This aligns with previous observations delineating a ‘mesenchymal signature’ as one of the major transcriptional programs present in primary cells from different tissue types, across developmental lineages^34^. Transitioning towards such a mesenchymal signature may transcriptionally prime different precursor lineages to increase their cell fate plasticity, akin to what is observed in various types of metastatic cancers, while still maintaining partially distinct transcription and chromatin memories of their embryonic origins^35,36^. Indeed, skeletogenic cells of the three lineages investigated here all start out as embryonic epithelia and undergo an epithelial-to-mesenchymal transition (EMT), before migrating to their respective anatomical locations in the periphery^37-39^. There, they form mesenchymal condensations and appear to initiate a core skeletogenic program which – at a global scale – shows similarity in its overall gene regulatory state^20,40,41^ (Fig.1H). We also noted a shared increase in ribosomal protein transcription, potentially to meet the translational needs for increased extracellular matrix protein production and secretion^42^ (Fig.S4A). Importantly, however, our cellularly resolved analyses along the mesenchymal precursor-to-skeletal progenitor trajectories also uncovered embryonic lineage-specific *trans*- and *cis*-regulatory dynamics during the formation of the vertebrate skeleton.

### Distinct *trans*- and *cis*-regulatory dynamics along skeletogenic differentiation trajectories of different embryonic origins

We find evidence for a lineage-specific heritage in transcription factor expression profiles that carries over to skeletogenic cells at different anatomical locations of the vertebrate skeleton (Fig.2B,C). This implies that the convergent specification of functionally analogous and transcriptionally similar skeletal cell types can be induced by distinct upstream *trans*-regulatory inputs, even across germ layers. Similarly distinct inputs with convergence were identified between different neuronal lineages in the developing visual system of *Drosophila*^43^. Here, using scATAC-seq data, we add an additional layer of regulatory information, and demonstrate that distinct chromatin accessibility signatures are accompanying these specific *trans*-regulatory inputs (Fig.2G,H). These lineage-specific *cis*-regulatory signatures separate unequally between gene-proximal and -distal sites, with promoter accessibilities exhibiting overall higher similarities in skeletal cells across embryonic origins than distal sites with putative enhancer functions (Fig.2I,J).

Using pseudotime analyses we followed the *trans*- and *cis*-regulatory dynamics along the skeletogenic differentiation trajectories of the three embryonic origins and identify both known and novel signatures of this transcriptional and phenotypic cell fate convergence (Fig.3D-I, Fig.S4A,D). Intriguingly, the expression levels of lineage-specific *trans* regulators along our pseudotemporal orderings peak right before the onset of the core skeletogenic program (Fig.3D-I), thus raising the possibility of lineage-specific transition states in which mesenchymal precursors are transcriptionally recoded towards a similar skeletogenic cell fate^44^. Thereafter, known skeletal regulators become transcribed in the respective convergence trajectories. We provide evidence that the activation of these genes relies on non-overlapping set of enhancer elements, with lineage-specific chromatin accessibility profiles and promoter-enhancer interactions (Fig.3J-L, Fig.5E,F). Using *in vivo* reporter assays, we demonstrate the lineage-specific activity of distal enhancer elements at a core chondrogenic factor (Fig.5G,H). Human genetics and molecular studies using *in vivo* and *in vitro* models have previously documented the presence of large regulatory landscapes at skeletogenic genes^45-47^. We argue that this regulatory strategy enables distinct up-stream *trans* inputs to activate a common downstream program in the different embryonic lineages, through integration at the level of *cis* elements (Fig.6). In this scenario, the implementation of distinct long-range enhancer actions would transcriptionally recode distinct precursor signatures towards a similar skeletogenic cell fate.

**Figure 6.**
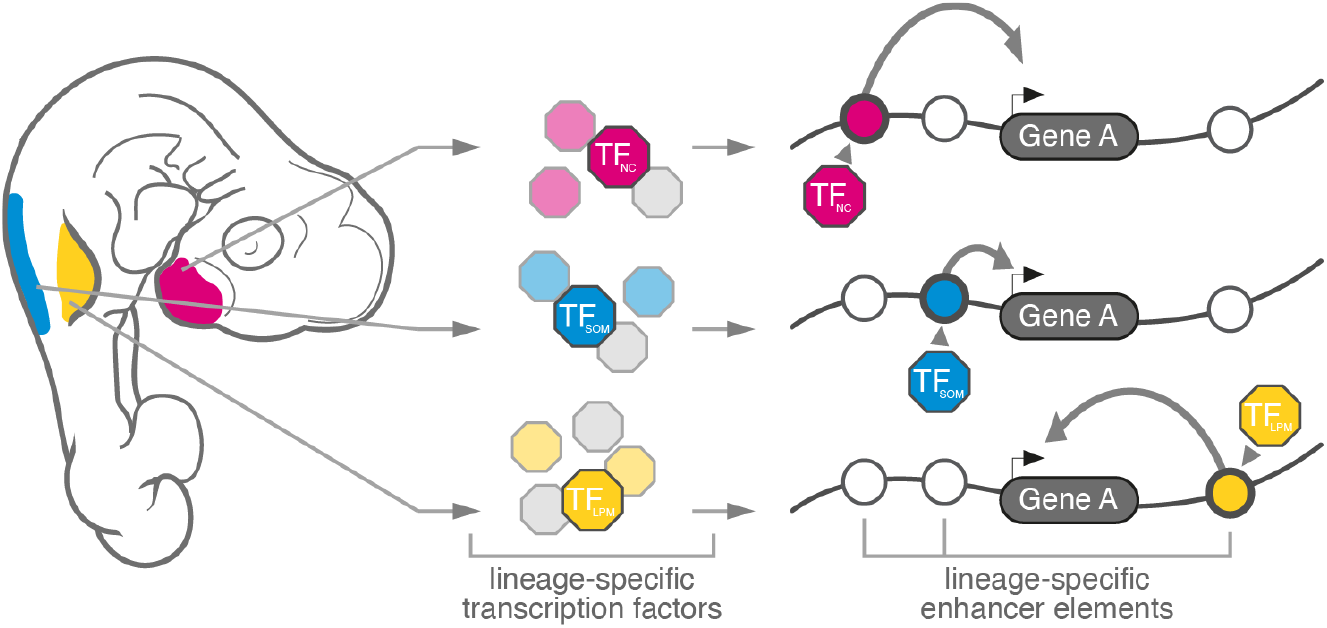
Model for the transcriptional convergence of skeletogenic cells of different embryonic origins. Depending on anatomical location, different embryonic lineages give rise to the skeletal elements of the vertebrate skeleton. These mesenchymal precursors express distinct lineage-specific transcription factor profiles, according to their embryonic origin, which are integrated at lineage-specific enhancer elements to activate a shared set of genes belonging to a core transcriptional program of vertebrate skeletal progenitors.

Across the different embryonic lineages, it is unlikely that this convergence involves only a single skeletogenic ‘master regulator’. Indeed, *SOX9* – a prime candidate for such function – acts in many non-skeletogenic lineages^48^, and its overexpression is unable to fully reprogram cells with skeletogenic potential towards a chondrocyte fate by itself^49^. Rather, an entire battery of transcriptional regulators – some shared, many lineage-specific – seems to drive the lineage-specific convergence towards a skeletogenic phenotype, with further distinctions present at the level of distal *cis* elements (Fig.3D-L, Fig.2H-J). In fact, at early stages of this convergence process, we find shared *trans*- and *cis*-evidence for only three previously identified factors, *SOX9, SOX5* and *FOXP1*^50^ (Fig.4F,G). This apparent regulatory complexity appears further potentiated by differences in transcription factor binding motifs, in mesenchymal cells across the embryonic origins (Fig.4H-J). Whether due to lineage-specific differences in local chromatin environment or the presence of distinct heteromeric binding partners, such expanded ‘regulatory grammar’ at the level of motif diversity is known to increase the combinatorial flexibility and complexity in cell fate decision and patterning processes^30-32^.

### Origin-specific evolutionary trajectories of skeletogenic cells

The distinct regulatory modalities that specify the skeletogenic cells in different parts of the vertebrate embryo also hold implications for how we treat them in an evolutionary comparative context. Indeed, thanks to ever more detailed molecular descriptions of cell types across the tree of life, the emerging ‘regulatory blueprints’ have helped to delineate what actually defines distinct cell types, and disentangle putative evolutionary histories and developmental relationships amongst them^1,2^. In vertebrates, different anatomical regions have acquired the potential to form an endoskeleton at distinct evolutionary timepoints. It is generally believed that an acellular, cartilage-like support of feeding structures preceded vertebrates^13^. A possible incorporation of the underlying gene regulatory network into skeletogenic precursor cells would then have paved the way for the emergence of the vertebrate cranium^51-53^. As additional developmental lineages gained skeletogenic competency, repeated co-option of such network would appear as the most parsimonious solution to developmentally specify a shared cellular phenotype from distinct embryonic sources. However, the partial regulatory independence we document here suggests network recruitment with modification, and thus only an incomplete ‘deep homology’ amongst early skeletogenic cells of different embryonic origins^17^. While these different regulatory strategies might originally have been a necessity – namely, to absorb and integrate the distinct *trans*- and *cis*-regulatory modalities of the evolutionarily novel skeletogenic precursor lineages – ultimately, they might have proven beneficial and even adaptive. By relying on partially distinct specification networks, independent changes in effector gene expression – e.g., to affect extracellular matrix composition and ensuing tissue properties^42,54^ – or cellular growth dynamics become possible in the different parts of the skeleton^55-57^. Indeed, naturally occurring genetic variation as well as induced targeted mutations in some of the embryonic origin-specific regulators we document here show anatomical location-restricted effects^57-59^. Hence, despite seemingly similar cellular phenotypes, the distinct regulatory strategies at work in the three embryonic lineages may help to make different parts of the vertebrate skeleton become independent targets of evolutionary selection, with distinct biomaterial and patterning properties resulting therefrom.

## Materials and Methods

### Tissue Collection

Embryos were dissected in ice-cold PBS and staged according to Hamburger and Hamilton^60^ (HH). Embryonic tissue was either processed for single-cell functional genomics experiments, or fixed in 4% PFA at 4C.

### RNA *In Situ* Hybridization

Cranial or brachial tissue samples ranging from HH17 to HH24 were dehydrated and cryo-embedded side-by-side in OCT, to allow for staining of entire time series on single slides. Sectioning was performed on a *Leica CM3050S* cryostat and RNA *in situ* hybridization against *SOX9* and *ACAN* was performed using standard protocols^61^. Brightfield images were acquired on an *Olympus FLUOVIEW FV3000* and globally processed for color balance and brightness using *Adobe Photoshop*.

### Single-Cell RNA-Sequencing (scRNA-seq) Data Collection

We sampled the frontonasal prominence at stages HH15, HH18 and HH22, the dorsal part of the brachial region at stages HH12, HH15 and HH20, and entire forelimbs at stages HH21, HH24 and HH27. Tissue dissociation, *10x Genomics Chromium 3’ Kit* library preparation and sequencing was performed as reported previously^21^. Data was processed and mapped with *CellRanger* (*10x Genomics*), using our in-house improved GRCg6a genome annotation with elongated 3’ UTRs^62^.

### scRNA-seq Data Processing

Unique Molecular Identifier (UMI) count matrices were filtered for quality based on a cell’s total and relative UMI counts (i.e., >4*mean and <0.2*median of the sample), percentage of mitochondrial UMIs (i.e., >median + 3*MAD (median absolute deviation) & >0.1, except if UMI count >median), and relation of UMI count/genes detected (i.e., <0.15 & UMI count <2/3) ^62^. In total, 8468 cells remained for frontonasal (2702, 4558 and 1208 cells for HH15, HH18 and HH22), 7777 cells for somite (3093, 2993 and 1691 cells for HH12, HH15 and HH20), and 10350 cells for forelimb (2987, 5293 and 2070 cells for HH21, HH24 and HH27).

### scRNA-seq Data Normalization, Dimensionality Reduction and Clustering

Using the *R* package *Seurat* (v4) ^19^, UMI counts were normalized by sequencing depth and log transformed. A cell cycle score was calculated using *SCRAN*^63^. Variations of sequencing depth, mitochondrial UMI percentage, and the difference in S and G2M cycle scores were regressed out, using *SCTransform* from *Seurat*^19,62,64^. Genes with a higher value of standardized variance than the sum of median and median absolute deviation were considered as ‘highly variable’. These steps were carried out independently for the three stages of the three embryonic origins.

Using *Seurat*, we integrated samples from the same embryonic origin, and used principal component analysis (PCA) on highly variable genes, followed by tSNE and FFT-accelerated Interpolation-based tSNE algorithms^65^ for nonlinear dimensionality reduction on the first 19 (nasal), 21 (somite) and 19 (limb) principal components. Using *Seurat* functions, we performed Leiden graph-based clustering^66^ on all cells with a resolution of 0.2 (=‘broad clustering’). A second round of clustering was conducted on select mesenchyme populations, with resolutions of 0.4 (somite, limb) and 0.5 (nasal) (=‘fine clustering’). Cell type assignments of clusters were based on visual inspection of known marker gene expression patterns, and the activity of the two previously identified early chondrogenic gene expression modules ‘IMM’^20^ and ‘RED’^21^ using the *Seurat* function ‘AddModuleScore’.

### Differential Expression Analysis

Differential expression analysis was based on a logistic regression framework^67^ using *Seurat*, with cell cycle differences and embryonic stages as latent variables. Genes expressed in at least 10% of the cells and showing differences with an adjusted p-value <0.05 and a log fold change >0.5 (‘broad’) or >0.25 (‘fine’) were considered as significantly differentially expressed. To minimize batch effects, differential expression analysis of chondrocytes from different embryonic origins was performed on pseudobulk counts using the *R* package *muscat*^68^.

### scRNA-seq Data Integration Across Embryonic Origins

We filtered out potential doublets using the *R* package *doubletFinder* ^69^ and removed clusters enriched for mitochondrial counts. The resulting UMI count matrix was divided by size factor and logtransformed using *SCRAN*^63^. The top variable genes (getTopHVGs, *SCRAN*) identified in at least two samples were kept for downstream analyses. Using *Seurat*, we then integrated the count matrices using anchors in canonical correlation analysis (CCA) reduction, to compute batch corrected matrices of the three embryonic origins. To calculate co-embedding projections, the PCA dimension were reduced sample-wide. Anchors for integration were identified using ‘FindIntegrationAnchors’ in reciprocal PCA reductions. We used ‘IntegrateEmbeddings’ to integrate PCA reduction, followed by tSNE calculations (‘RunTSNE’). Correlation analyses were performed on ‘pseudobulk’ average gene expression values (*Seurat* function ‘AverageExpression’) in each cluster.

### scRNA-seq Pseudotime Analyses

We generated spliced/unspliced count matrices of our selected mesenchymal populations using *velocyto*^70^ and assessed the directional transcriptional dynamics of highly variable genes with sufficient spliced/unspliced counts in *scVelo* with the default parameters^22^. We visualized the recovered dynamics on tSNE projections of the three embryonic origins^65^. We then used these tSNE embeddings as input space, and constructed a minimum spanning tree with preset start cluster in the *R* package *slingshot*^25^. Alignment of embryonic origin-specific pseudotimes was performed with *TrAGEDy*^26^ using 40 interpolated points along the respective chondrogenic trajectories. Module expression dissimilarities were calculated by Spearman correlation (1-ρ) and optimal alignment was identified by dynamic time warping with default settings. Using the *R* package *tradeSeq*^27^, we detected temporally differentially expressed genes along the respective chondrogenic trajectories.

### Single-Cell ATAC-Sequencing (scATAC-seq) Data Collection

Tissue dissociation was performed as previously described^21^. Cell concentration and viability were assessed using the *Nexcelom Cellometer K2* and ∼1*10^6^ cells were used to perform nuclei isolation following the *10x Genomics* protocol. Nuclei suspensions were loaded onto a Next GEM chip H, and transposition, nuclei partitioning and library preparation were performed according to the *10X Genomics* ATAC User Guide. scATAC libraries were quantified on an Agilent 2100 Bioanalyzer system (Agilent) and sequenced on a NovaSeq 6000 system (Illumina).

We used *CellRanger ATAC v1*.*2*.*0* (*10x Genomics*) for read processing and quantification, and mapped the fragments to the chicken ENSEMBL genome Gallus_gallus-6.0^71^ with our in-house improved GRCg6a genome annotation^62^.

PEAK_MERGE_DISTANCE was changed to 50, with all other parameters at default settings. **scATAC-seq Data Pre-Processing**. We removed doublets with *ArchR* (v1.0.1) ^33^ and selected high quality cells in *Signac* (v1.1.1) ^72^ using the following thresholds: total number of fragments in peaks ranging from 1000 to 100000, fraction of reads in peaks >15%, nucleosome signal < 4 and TSS enrichment score > 2. Using these criteria, we ended up with 11527 cells for frontonasal (6171 and 5356 cells for HH15 and HH18), 14106 cells for somite (11232 and 2874 cells for HH12 and HH15), and 10982 cells for forelimb (4453 and 6529 cells for HH21 and HH24).

### scATAC-seq Data Merging, Dimensionality Reduction and Clustering

We merged samples from same embryonic origins and summed fragment counts in 5kb genomic tiling windows located on autosomes and chromosome Z (208680 tiles in total). We performed latent semantic indexing (LSI) dimension reduction on a term frequency-inverse document frequency (TF-IDF) normalized matrix with the top 75% of tiles (top 0.1% tiles are removed, putative repetitive elements or alignment errors) using *Signac* and removed batch effects on LSI components using *Harmony*^73^. We used tSNE and FFT-accelerated Interpolation-based t-SNE algorithm^65^ to carry out non-linear dimensionality reduction with LSI dimensions 2:30, and performed Leiden graph-based clustering in *Seurat* on all cells with resolutions of 0.4 (nasal, somite) and 0.6 (limb) (=‘broad clustering’), and a second round of clustering on select mesenchyme populations with resolution of 0.4 (=‘fine clustering’). We annotated cell types for both ‘broad’ and ‘fine’ clusters using scATAC-seq gene activity matrices (promoters and gene bodies) and scRNA-seq expression data, with a combination of label transfers in *Seurat* and non-negative least squares (NNLS) regression on cluster specific genes^74^, as well as manual inspection of peaks at known marker genes.

### Peak Calling and Differential Accessibility Analysis

We identified peaks using *MACS2* (version 2.2.7.1) ^24^ with parameters “—nomodel --shift 100 --extsize 200 --keep-dup all --call-summits” on pseudobulks of each cluster, for each embryonic origin, respectively. Peaks used summits as center and were extended to a width of 501bp. We merged peaks from different clusters of the same embryonic origin and, for overlapping peaks, kept only the most significant one, using adapted code from *ArchR*. To get a consensus peak set, we merged peaks from three embryonic origins and removed redundant and/or overlapping peaks using the same logic.

Differential accessibility analysis was performed in *Seurat*, using the total number of fragments and embryonic stages as latent variables. Peaks accessed in at least 10% of the cells and showing differences with adjusted p-value less than 0.05 and a log fold change larger than 0.25 were considered as significantly differentially accessed. Peak-centered heatmaps of differential accessible peaks were visualized with *deepTools2*^75^.

### scATAC-seq Data Integration Across Embryonic Origins

The top variable peaks (getTopHVGs, *SCRAN*) identified in at least two samples were kept for downstream analyses. To calculate a co-embedding projection, first we performed reciprocal LSI dimensional reduction to find anchors (*Seurat* function ‘FindIntegrationAnchors’) and constructed transformation matrices between each query cell and anchor. We computed the integration matrices based on the original LSI matrix with dimensions from 2 to 30 and the transformation matrix using the *Seurat* functions ‘IntegrateEmbeddings’ and ‘runTSNE’ on the integrated LSI dimensions. To remove the batch effects among peak matrices after merging, we binarized the matrix based on presence/absence of counts. Correlation analyses were performed on ‘pseudobulk’ average count values in each cluster using the function ‘AverageExpression’.

### scATAC-seq Pseudotime Analyses

We transferred pseudotime values from our scRNA analyses using the *Seurat* function ‘TransferData’. First, we integrated scRNA-seq expression matrices and scATAC-seq gene activity matrices and performed CCA dimensional reduction to find anchors. We then constructed a transformation matrix between each query cell and each ancho (*Seurat* function ‘FindTransferAnchors’) and computed the transferred scATAC-seq pseudotimes based on the original scRNA-seq pseudotimes and the transformation matrices. For all three embryonic origins, we restricted this transfer to only chondrogenesis related cell type clusters.

### *de novo* Motif Enrichment Analysis and Annotation

We performed *de novo* motif enrichment analysis for each cluster, using *Homer*^28^ ‘findMotifsGenome.pl’ with -mset vertebrates -size -250,250 -fdr 5 and motif length between 8-22 bp (*Homer* p-value < 1e-11), using highly accessible peaks for each cluster. We obtained candidate TF annotations for this set of *de novo* motifs with the help of three databases (*Homer* vertebrates, *JASPAR20* vertebrates^76^, *CisBP* v2 chicken^77^) using *Homer* and *STAMP*^78^. We selected the best matches based on scRNA-seq expression levels of the predicted TFs, and Spearman correlations between motif activity and gene expression of candidate TFs in scATAC and scRNA aggregates (default k=50, n=400; for small clusters, k=20, n=200). For motifs with similarity scores >0.8, only the one with the lowest p-value was retained. Additionally, we checked for paralog TFs expression along our pseudotime trajectories and calculated its expression correlation to motif activity. We combined annotated *de novo* motifs from each embryonic origin and calculated motif similarity scores using *PWMEnrich*^79^. For our final set of annotated *de novo* motifs, we computed a per-cell motif deviation scores using *chromVAR*^80^ and conducted analysis of differential motif activity using *Seurat*.

### Peak-to-Gene Link Analysis

We generated imputed pseudoexpression data for each scATAC cell based on scRNA-seq data, using the *ArchR* function ‘addGeneIntegrationMatrix’. 500 cell aggregates were generated with, with 100 cells per aggregate. We then computed the Pearson correlation between peak accessibility and pseudoexpression in mesenchymal aggregates using the *ArchR* function ‘addPeak2GeneLinks’. Clustering of peak-to-gene links was calculated by the hkmeans method in *factoextra*^81^. Functional enrichment analyses of peak-to-gene link clusters was conducted in rGREAT^82^.

### Enhancer Reporter Assays

Genomic regions of candidate enhancers were amplified by PCR from chicken genomic DNA and cloned upstream of a minimal promoter driving GFP expression. Fertilized chicken eggs were incubated at 38.5 °C in a humified incubator to stage HH14 for lateral plate mesoderm and somite electroporation and stage HH7 for neural crest electroporation. DNA solutions containing our enhancer reporter constructs and a constitutively expressing tdTomato co-electroporation control were injected and electroporated into epithelial precursor populations of the cranial neural crest, the brachial somites or the lateral plate mesoderm at forelimb levels^39,83,84^. Embryos were harvested two days post-electroporation and tissue was processed for immunohistochemistry^85^.

## Author Contributions

This study was conceived and designed by MW, ADPT, CF and PT. Single-cell functional genomics data was generated by CF, ML, CM and AF. Bioinformatics analyses were conducted by MW, CF and PT. Enhancer reporter assays were performed by ADPT and SF. PT wrote the paper, with feedback from all other authors.

## Acknowledgments

The authors wish to thank I. Adameyko and all members of the lab for insightful discussions. Calculations for single-cell functional genomics analyses were performed at sciCORE (http://scicore.unibas.ch/), scientific computing center at the University of Basel. This work was supported by research funds from the Swiss National Science Foundation [SNSF project grant number 310030_189242], the Swiss 3R Competence Centre [3RCC grant OC-2018-005] and the University of Basel to PT.

## Supplementary Figures

**Figure S1.**
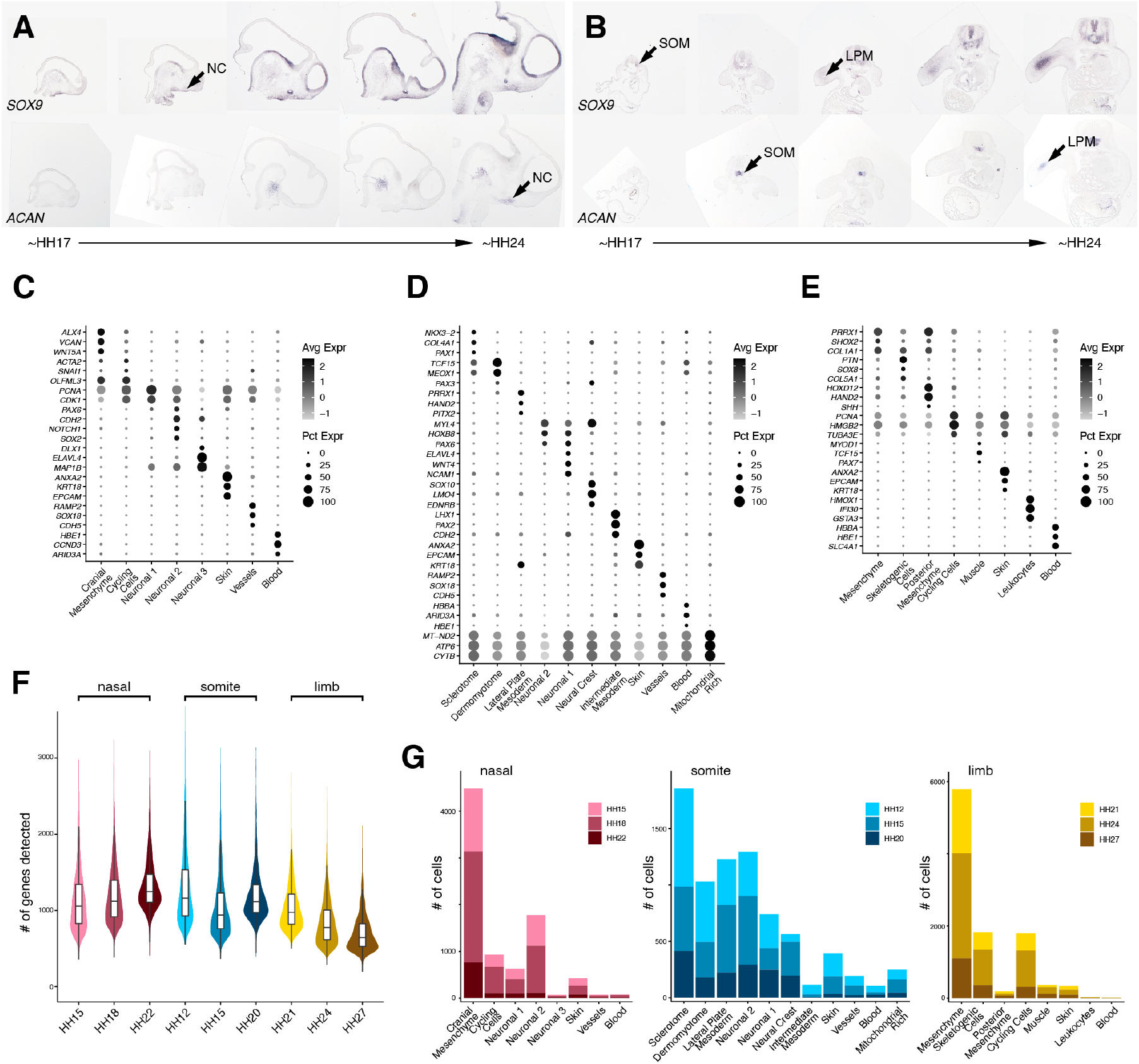
*In situ* dynamics of skeletogenic induction across three embryonic lineages and quality metrics of single-cell transcriptomics experiments. (**A, B**) Chromogenic *in situ* hybridizations against *SOX9* and *ACAN* on cranial sagittal (**A**) and brachial transversal (**B**) cryosections spanning embryonic stages HH17 to HH24. First appearance of signal in the respective embryonic lineages is marked by an arrow (NC: neural crest, SOM: somite, LPM: lateral plate mesoderm). (**C**-**E**) Dot plots of select marker genes expression used for ‘broad’ cluster cell type annotations. (**F**) Violin plots of number of genes detected in the nine scRNA-seq samples, color coded by anatomical origin and embryonic stage. (**G**) Stacked bar plots indicating stage-wise contributions to the total number of cells in ‘broad’ clusters, color coded by anatomical origin and embryonic stage.

**Figure S2.**
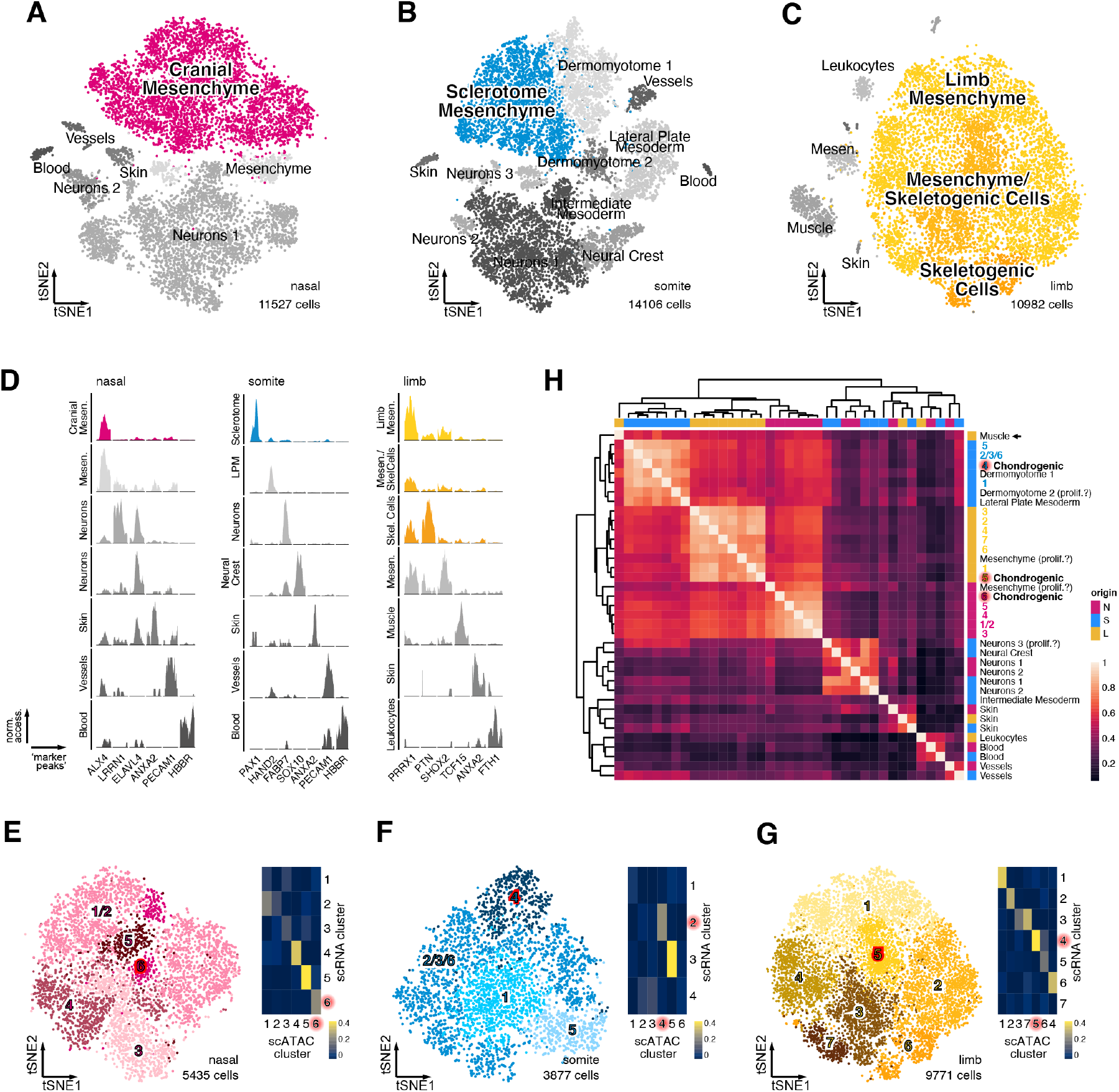
The single-cell chromatin accessibility landscape of skeletogenic induction across three embryonic lineages. (**A**-**C**) tSNE representations of the three scATAC-seq datasets and ‘broad’ cell type annotations, with mesenchymal populations used for ‘fine’ re-clustering highlighted in color. (**D**) Pseudobulk aggregate genome accessibility tracks of peaks near ‘broad’ cell type-specific marker genes. (**E**-**G**) tSNE representations of re-clustered mesenchymal cells of nasal (**E**), somite (**F**) and limb (**G**) origins, with ‘fine’ cluster annotations and normalized scores based on non-negative least squares (NNLS) regression, between gene activity from ATAC clusters and gene expression from RNA clusters. High values are indicative of cell type correspondences between scATAC and scRNA clusters. Early chondrogenic clusters are highlighted in red. (**H**) Unsupervised hierarchical clustering and heatmap representation of pairwise Spearman’s rank correlation coefficients of differentially accessible peaks in pseudobulk chromatin accessibility data of ‘broad’ and ‘fine’ mesenchymal clusters. Anatomical origins of pseudobulks are indicated by color code. All mesenchymal pseudobulks cluster by embryonic origins, including early chondrogenic cells (highlighted in red) and somite-derived muscles in the limb sample (arrow).

**Figure S3.**
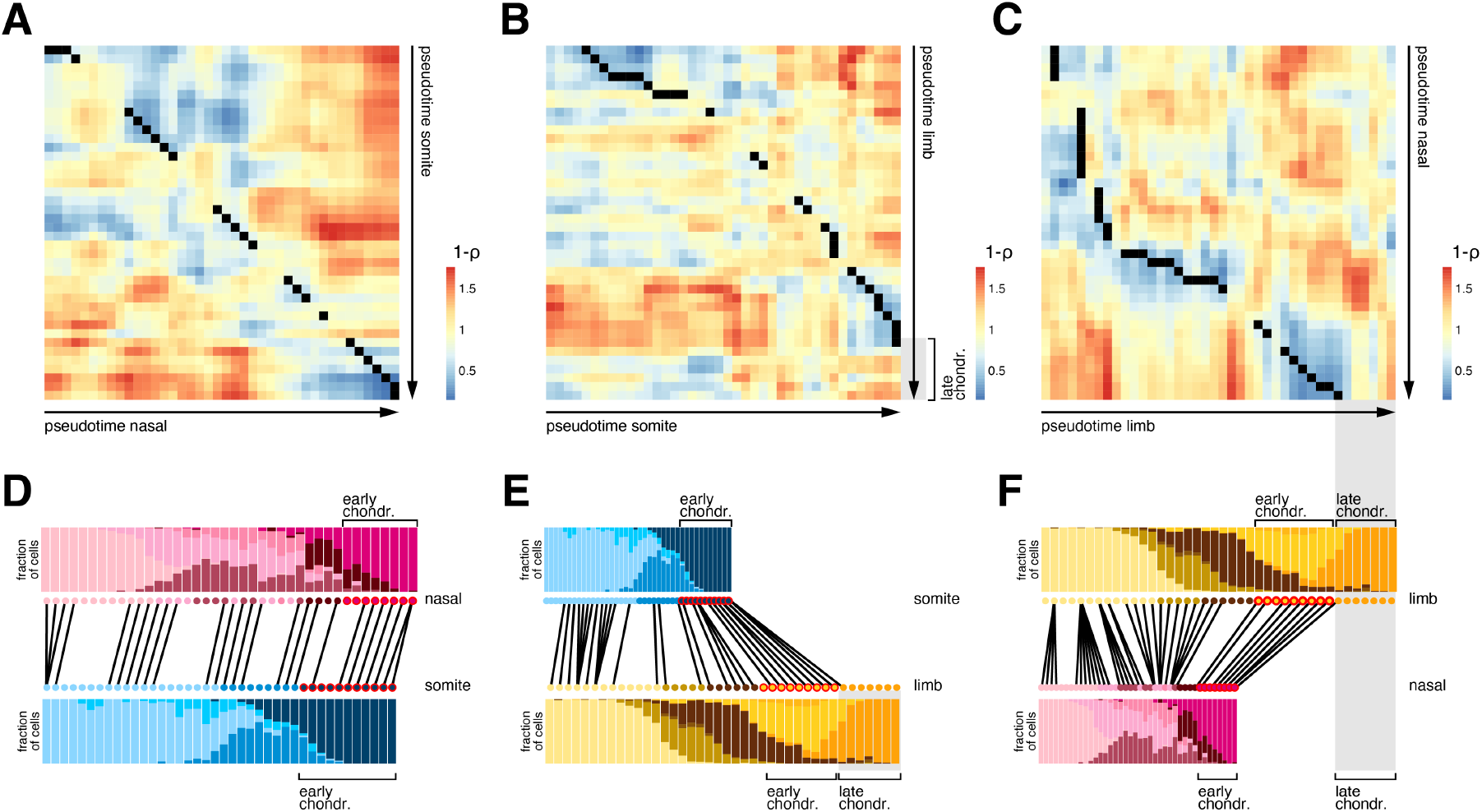
Skeletogenic convergence across three embryonic lineages. (**A**-**F**) *TrAGEDy* alignments of embryonic origin-specific chondrogenic pseudotime trajectories. (**A**-**C**) Pairwise expression dissimilarities of the chondrogenic module ‘IMM’ were calculated by Spearman correlation across all pseudotimes and embryonic origins, and are displayed as 1-ρ. Dynamic time warping identifies skeletogenic convergence across all three embryonic lineages, except for the ‘late chondrocytes’ population in the limb sample that is not accounted for in our nasal and somite data. (**D**-**F**) *TrAGEDy* alignment plots of interpolated points of chondrogenesis trajectories and cell type contributions per pseudotime bin, displayed as stacked bar plots. Pseudotime bins where most cells are classified as ‘early chondrogenic’ are highlighted in red.

**Figure S4.**
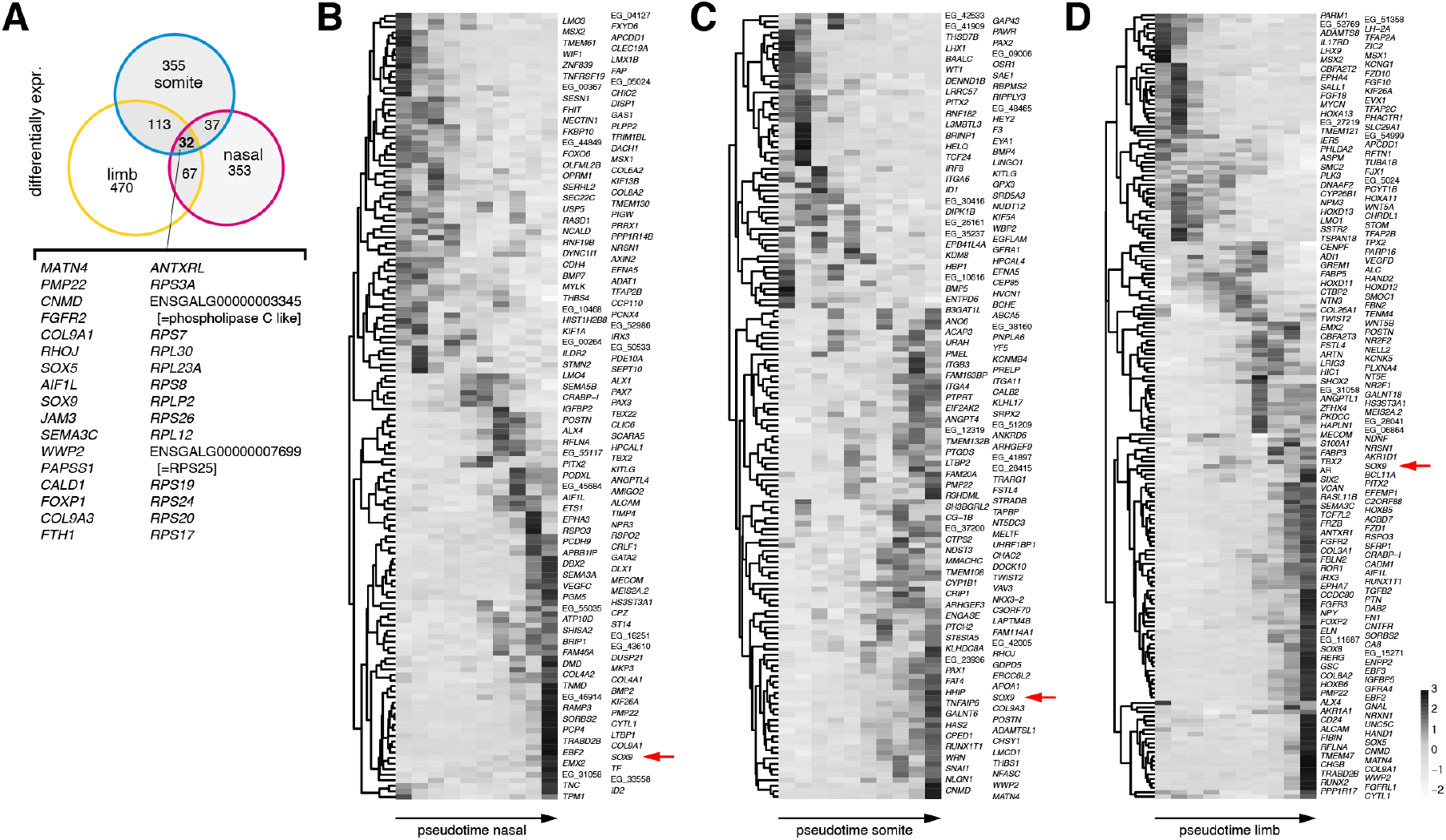
Core skeletogenic genes and embryonic origin-specific transcriptional dynamics. (**A**) Venn diagram displaying the overlap of differentially expressed genes in chondrogenic cells compared to other mesenchymal populations of the three embryonic origins. (**B**-**D**) Z-score scaled RNA levels of dynamically expressed genes along the chondrogenic trajectories of nasal (**B**), somite (**C**) and limb (**D**) mesenchymal and chondrogenic cells. *SOX9* expression is highlighted by a red arrow. Genes without annotated symbol are labeled with ‘EG_’, followed by the 5 last digits of their ENSGAL identifiers.

**Figure S5.**
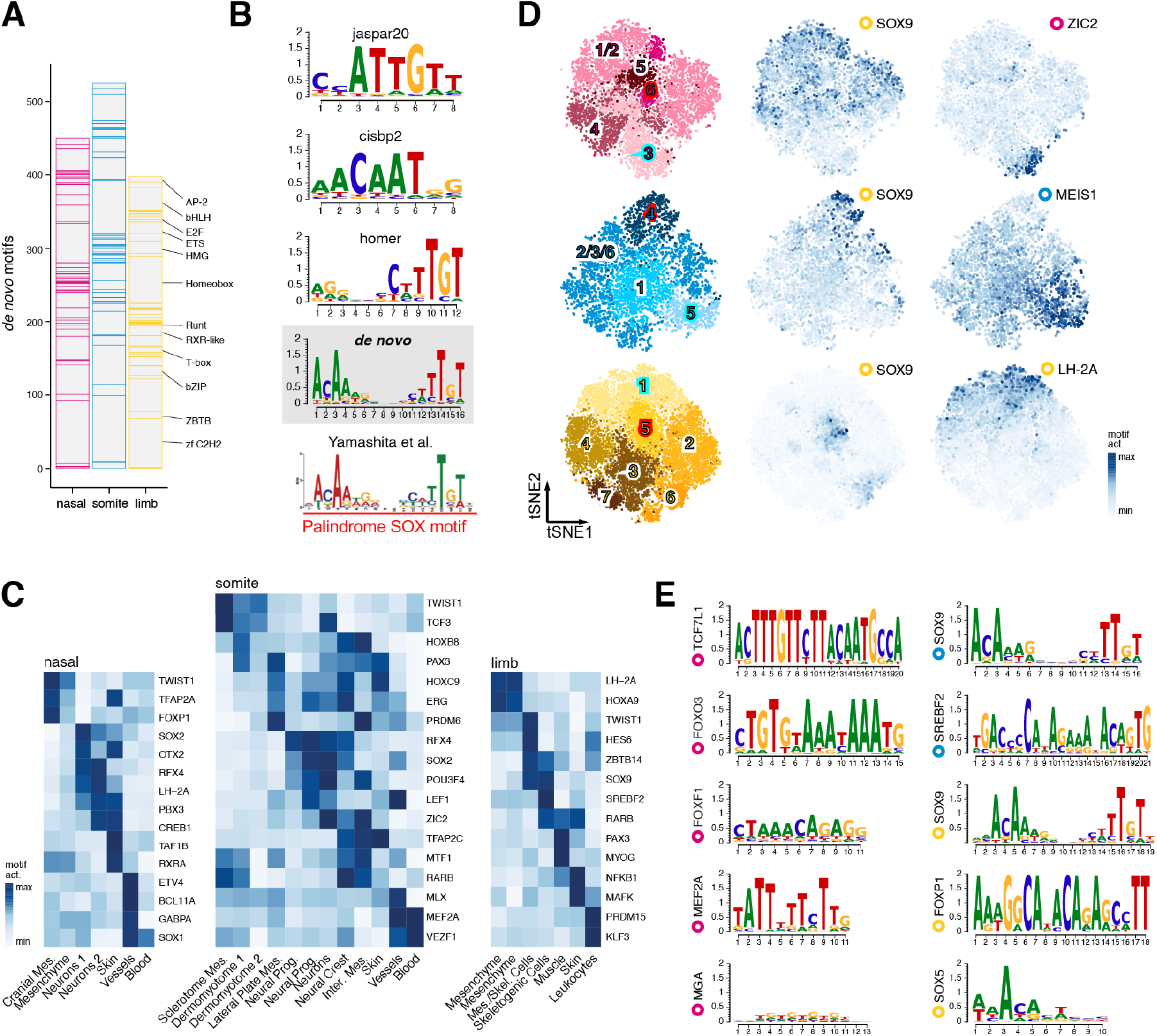
*De novo* identification and annotation of transcription factor binding motifs. (**A**) Transcription factor (TF) family distribution of all *de novo* identified binding motifs in all populations across the three embryonic origins. (**B**) Publicly available position weight matrices for SOX9 in ‘jaspar20’, ‘cisbp2’ and ‘homer’ databases. Our *de novo* position weight matrix resembles more closely the one identified using SOX9 ChIP-seq data in chicken embryonic limbs^1^. (**C**) Differential ‘broad’ cluster-specific motif activities across embryonic origins. (**D**) tSNE representations of reclustered mesenchymal cells, with motif activity heatmaps plotted for a core skeletogenic factor (SOX9, middle) and an embryonic origin-specific precursor factor (ZIC2, MEIS1, LH-2A, right). ‘Early chondrogenic’ clusters are highlighted in red, ‘mesenchymal’ clusters in turquoise (left). (**E**) Position weight matrices of the ten commonly chondrocyte-enriched TF motifs across embryonic origins. The embryonic origin in which the respective motif was identified is indicated by color-coded circles.

## Footnotes

1 SOX9 ChIP-seq position weight matrix from Yamashita et al.^29^, re-used under CC-BY 4.0 license

